# The nuclear localization sequence mediates hnRNPA1 amyloid fibril formation revealed by cryoEM structure

**DOI:** 10.1101/2020.06.05.135673

**Authors:** Yunpeng Sun, Kun Zhao, Wencheng Xia, Jinge Gu, Yeyang Ma, Xinrui Gui, Xia Zhang, Yanshan Fang, Bo Sun, Cong Liu, Dan Li

**Affiliations:** Interdisciplinary Research Center on Biology and Chemistry, Shanghai Institute of Organic Chemistry, Chinese Academy of Sciences, Shanghai 201210, China; University of Chinese Academy of Sciences, Beijing 100049, China; School of Life Science and Technology, ShanghaiTech University, Shanghai, 201210, China; Bio-X-Renji Hospital Research Center, Renji Hospital, School of Medicine, Shanghai Jiao Tong University, Shanghai, 200240, China; Bio-X Institutes, Key Laboratory for the Genetics of Developmental and Neuropsychiatric Disorders, Ministry of Education, Shanghai Jiao Tong University, Shanghai, 200030, China

## Abstract

Human heterogeneous nuclear ribonucleoprotein A1 (hnRNPA1) serves as a key regulating protein in RNA metabolism. Malfunction of hnRNPA1 in nucleo-cytoplasmic transport or dynamic phase separation leads to abnormal amyloid aggregation and neurodegeneration. The low complexity (LC) domain of hnRNPA1 drives both dynamic phase separation and amyloid aggregation. Here, we use cryo-electron microscopy to determine the amyloid fibril structure formed by hnRNPA1 LC domain. Remarkably, the structure reveals that the nuclear localization sequence of hnRNPA1 (termed PY-NLS), which is initially known to mediate the nucleo-cytoplamic transport of hnRNPA1 through binding with karyopherin-β2 (Kapβ2), represents the major component of the fibril core. The residues that contribute to the binding of PY-NLS with Kapβ2 also exert key molecular interactions to stabilize the fibril structure. Notably, hnRNPA1 mutations found in familial amyotrophic lateral sclerosis (ALS) and multisystem proteinopathoy (MSP) are all involved in the fibril core and contribute to fibril stability. Our work illuminate structural understandings on the pathological amyloid aggregation of hnRNPA1 and the amyloid disaggregase activity of Kapβ2, and highlights the multiple roles of PY-NLS in hnRNPA1 homeostasis.

## Introduction

Heterogeneous nuclear ribonucleoprotein A1 (hnRNPA1) is a multifunctional protein playing key roles in the regulation of gene expression and RNA metabolism^1–3^. It is involved in a wide-range of RNA processing events under normal and pathological cellular conditions including transcription, constitutive and alternative splicing, stability, nuclear export and translation^4–7^. HnRNPA1 contains two globular RNA recognition motifs (RRM1 and RRM2) for the specific binding with mRNA precursor, followed by an unstructured low complexity (LC) C-terminal domain. HnRNPA1 LC consists of a cluster of RGG repeats with interspersed aromatic residues and a nuclear localization sequence (termed PY-NLS)^8–10^. HnRNPA1 LC is intrinsically disordered and exhibits a characteristic feature of homogenous and heterogeneous supermolecular polymerization^11–14^. It can mediate protein-protein and protein-RNA interactions to undergo liquid-liquid phase separation to participate in stress granule assembly ^11,12.^ The LC domain also exerts a prion-like function to mediate the amyloid aggregation of hnRNPA1, which is causative to neurodegenerative diseases including amyotrophic lateral sclerosis (ALS) and multisystem proteinopathy (MSP)^7,12,15–17^, and enhanced by inheritable mutations^12,15.^

HnRNPA1 predominantly localizes in nucleus, while shuttles between the nucleus and cytoplasm in response to cellular signals^11,18,19^. The transport of hnRNPA1 between nucleus and cytoplasm is mediated by karyopherin-β2 (Kapβ2, also known as transportin-1) via direct binding of the PY-NLS sequence with Kapβ2^2,9,10^. The complex structure of Kapβ2 and PY-NLS reveals a high-affinity binding of PY-NLS in the C-terminal arch of Kapβ2^8^. Moreover, recent studies demonstrate that Kapβ2 engages PY-NLS or analogous NLSs to inhibit and even reverse the amyloid fibril formation of hnRNPA1 and other RNA-binding proteins^20^.

In this work, we determine the atomic structure of amyloid fibril formed by hnRNPA1 LC domain by cryo-electron microscopy (cryoEM). The structure reveals that the fibril core is mainly formed and stabilized by PY-NLS. Mutations of hnRNPA1 identified in familial ALS and MSP patients are all involved in the fibril core and contribute to fibril stability. This structure not only demonstrates the role of PY-NLS in driving hnRNPA1 amyloid aggregation at the atomic level, and also renders mechanistic understanding for the anti-amyloid activity of Kapβ2.

## Results

### CryoEM structure determination of the amyloid fibril formed by hnRNPA1 LC domain

Recombinant hnRNPA1 LC (residues 186-320) was incubated at 16 °C for 2 days to form amyloid fibrils (**Fig. 1a**). The fibrils were fixed on a carbon grid and frozen in liquid ethane. CryoEM data were collected by using a 300 keV Titan Krios microscope. 62,656 fibrils from 3,227 micrographs were selected for the reconstruction of hnRNPA1 LC fibril (**Table 1**). 2D classification showed one dominant species of left-handed twist fibril (~97 % of the selected fibrils) (**Supplementary Fig. 1**). We then performed helical reconstruction by Relion^21^ and obtained the 3D density map of the hnRNPA1 LC fibril to an overall resolution of 2.8 Å (**Fig. 1b and Supplementary Fig. 2**). The density map showed two protofilaments intertwining along an approximate 2-fold screw axis to form a left-handed helix with a width of ~7 nm and a half pitch of ~ 45 nm (**Fig. 1c**). The helical twist and rise between neighboring hnRNPA1 LC subunits are 179.05° and 2.37 Å, respectively (**Fig. 1c and Table 1**). Based on the high-resolution cryoEM density map, we were able to unambiguously build an atomic structure model for hnRNPA1 LC fibril (**Fig. 1d**). Despite that the fibrils were formed by hnRNPA1 residues 186-320, only residues 251-295 were involved in the fibril core (FC) (**Fig. 1d**).

**Table 1.** Statistics of cryoEM data collection and refinement.

|  |  |
| --- | --- |
| Name | **hnRNPA1 LC fibril** |
| PDB ID | 7BX7 |
| EMDB ID | EMD-30235 |
| **Data Collection** |  |
| Magnification | 130,000 |
| Pixel size (Å) | 0.52 |
| Defocus Range (µm) | -1.3 to -2.1 |
| Voltage (kV) | 300 |
| Camera | K2 summit |
| Microscope | Titan Krios |
| Exposure time (s/frame) | 0.25 |
| Number of frames | 32 |
| Total dose (e<sup>-</sup>/Å<sup>2</sup>) | 59 |
| **Reconstruction** |  |
| Micrographs | 3,227 |
| Manually picked fibrils | 62,656 |
| Box size (pixel) | 288 |
| Inter-box distance (Å) | 30.0 |
| Segments extracted | 801,707 |
| Segments after Class2D | 259,615 |
| Segments after Class3D | 69,037 |
| Resolution (Å) | 2.80 |
| Map sharpening B-factor (Å<sup>2</sup>) | -109.473 |
| Helical rise (Å) | 2.37 |
| Helical twist (°) | 179.05 |
| **Atomic model** |  |
| Non-hydrogen atoms | 1,896 |
| Protein residues | 270 |
| Ligands | 0 |
| r.m.s.d. Bond lengths | 0.009 |
| r.m.s.d. Bond angles | 0.833 |
| All-atom clash score | 15.34 |
| Rotamer outliers | 0.00% |
| Ramachandran Outliers | 0.00% |
| Ramachandran Allowed | 9.30% |
| Ramachandran Favored | 90.70% |

**Figure 1.**
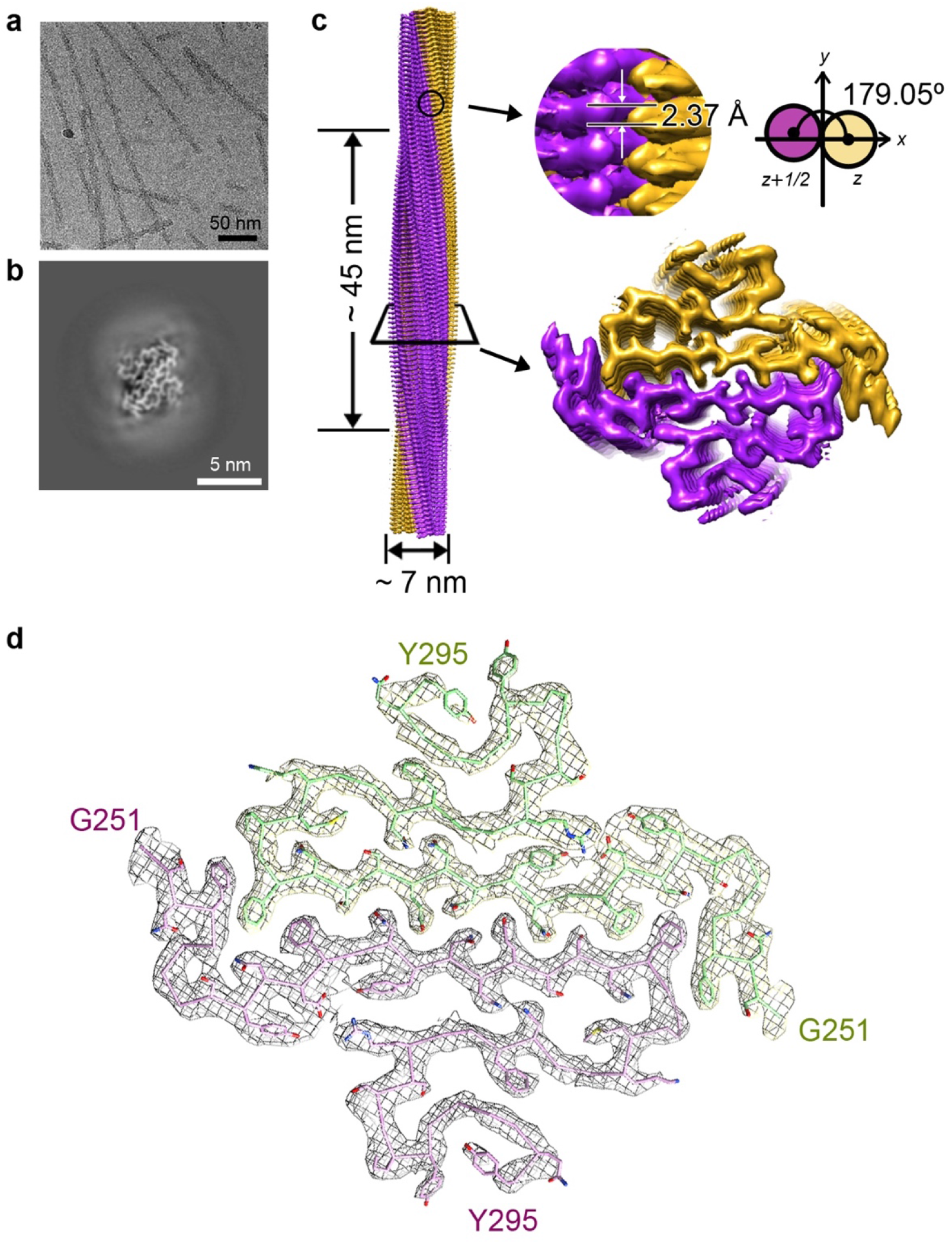
CryoEM structure determination of the amyloid fibril formed by hnRNPA1 LC domain. **a**, CryoEM image of the amyloid fibrils formed by hnRNPA1 LC. **b**, CryoEM 3D reconstruction of the hnRNPA1 LC fibril. **c**, CryoEM density map of the hnRNPA1 LC fibril. Fibril parameters including fibril width, half pitch, twist angle and rise are marked. Protofilaments are colored in magenta and orange individually. **d**, Cross-section view of the density map with structure model built upon it. Protofilaments are in different colors.

### Overall structural features of the hnRNPA1 LC fibril

HnRNPA1 LC is rich in glycine residues dispersed throughout the sequence. This property is reflected in a high content of random coils in the fibril structure (**Fig. 2a and 2b**). Although the coil regions lack rigid secondary structures, they orderly stack along the fibril axis to enhance the stability of each protofilament (**Fig. 2b and 2c**). In particular, coil 2 at the C terminus swaps to the next subunit to enhance the intra-protofilamental stability (**Fig. 2c**); coils 1 and 2 form inter-protofilamental contacts further enhancing the stability of the whole fibril (**Fig. 2b and 2c**).

**Figure 2.**
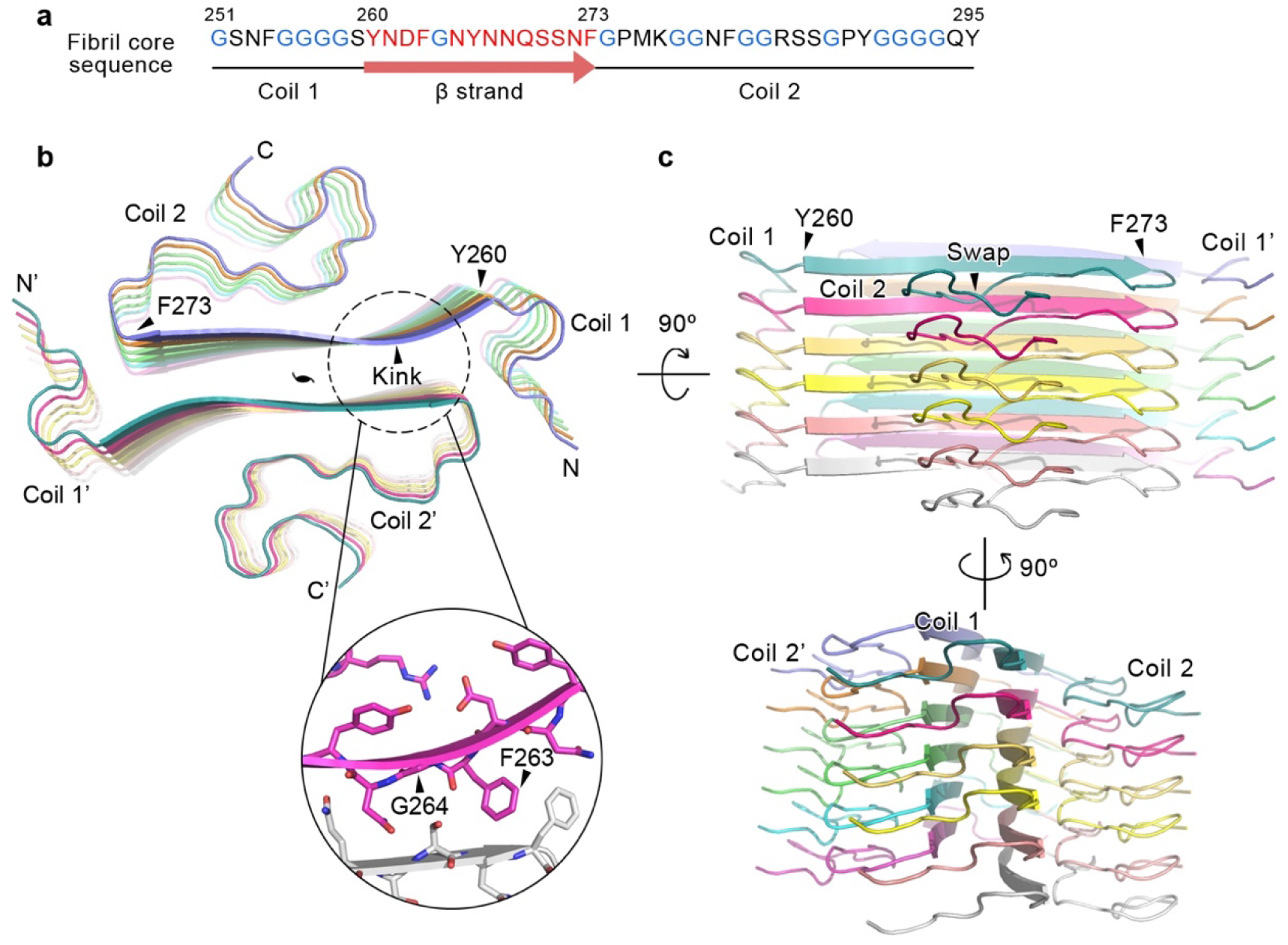
Overall structure of the hnRNPA1 LC fibril. **a**, The primary sequence of the hnRNPA1 FC. Residues forming the β strand are highlighted in red. Glycine residues are in blue. Secondary structures are indicated below. **b**, Top view of the cryoEM structure of hnRNPA1 LC fibril. Six layers of backbones are shown. The structure is colored by chain. The pseudo-2-fold screw axis is indicated. The sheet kink region is zoomed in with side chains shown. **c**, Side views of the fibril structure in (**b**).

The serpentine fold of hnRNPA1 FC remarkably contains a single but long β strand composed of residues 260-273 (**Fig. 2a and 2b**). β strands of the same protofilament form a parallel β sheet with a kink at _263_FG_264_ (**Fig. 2b**). A pair of β sheets mate to provide major inter-protofilamental interactions for the fibril formation (**Fig. 2b**). Of note, ordered coil and kinked β sheet structures have also been found in the reversible fibrils formed by FUS and hnRNPA1 segments^12,22–24^. Thus, these structural features might be general in Gly-rich LC domains.

### The hnRNPA1 LC fibril features a remarkably large protofilamental interface

Based on the fibril structure, residues 254-275, which represent half of the FC, are involved in the protofilamental interface (**Fig. 3a and 3b**). The interface renders a large buried surface area of 1,814 Å^2^ per layer. According to the local resolution estimation, the cryoEM density in this area is of the highest resolution of the entire fibril structure indicating that the protofilamental interface is the most stable region of the fibril (**Supplementary Fig. 2b**). The interface is tightly packed with a shape complementarity (Sc)^25^ of 0.844 excluding any cofactor or water molecules. It is formed by a typical parallel in-register steric-zipper in the middle sandwiched by Phe clusters (**Fig. 3b**). In the steric zipper, the opposing polar side chains of N265, N267, Q269, and S271 are tightly interdigitated to each other via van der Waals forces to form a shape complimentary dry interface (**Fig. 3c**), which is reminiscent of the first atomic structure of steric zipper formed by segment GNNQQNY of yeast prion Sup35^26^ (**Supplementary Fig. 3**). Flanking this steric zipper, F263 and the opposing F273 form T-shaped π stacking and further stack along the fibril axis (**Fig. 3a, 3b and 3d**). Outside the π stacking, F254 and the opposing P275 form hydrophobic interactions, which further enlarge the protofilamental interface, and together with the π stacking of F263-F273, stabilize the fibril by blocking the contact of solvent with the hydrophilic steric-zipper interface (**Fig. 3a, 3b and 3e**).

**Figure 3.**
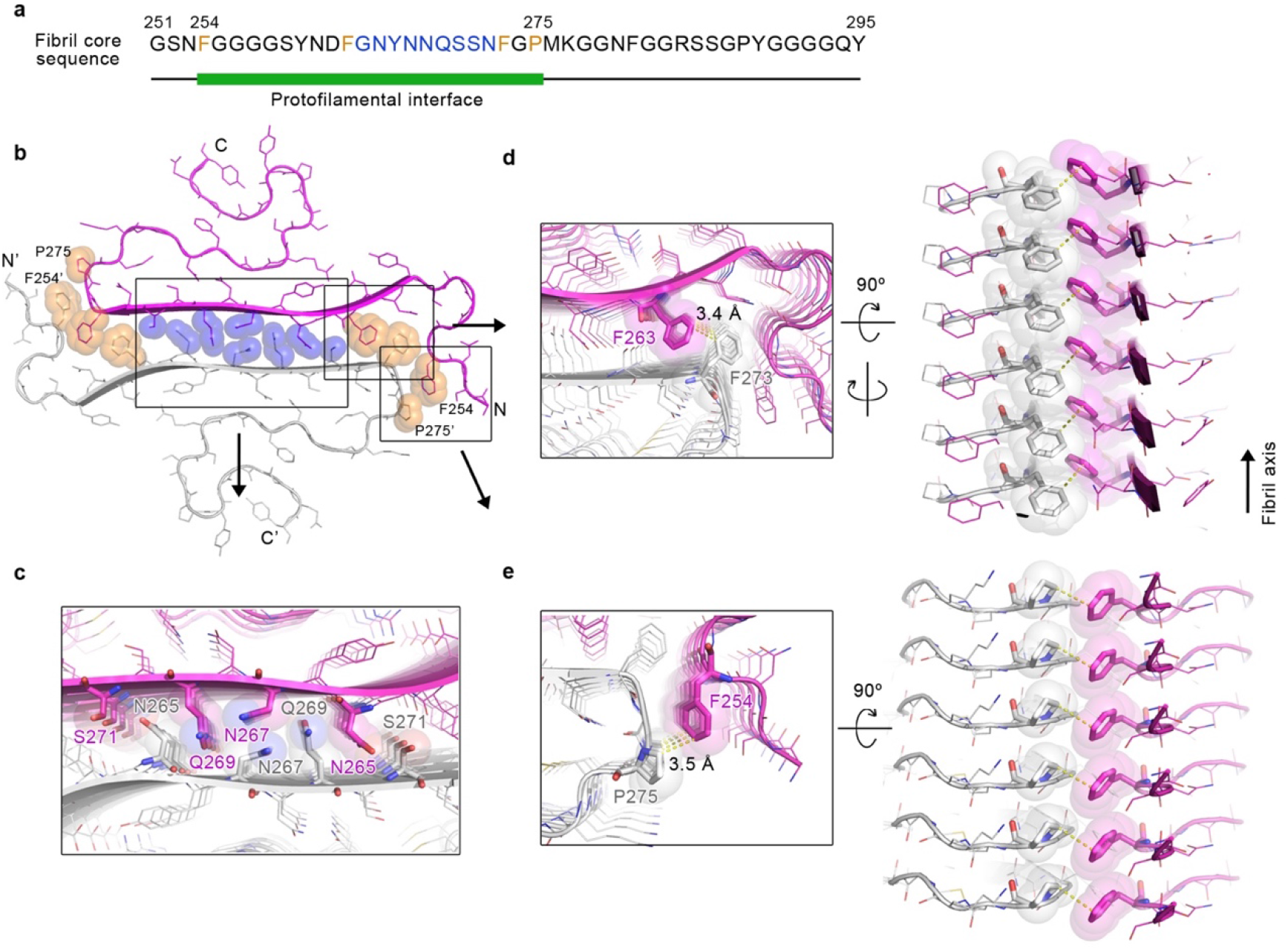
Protofilamental interface of the hnRNPA1 LC fibril. **a**, The primary sequence of the hnRNPA1 FC. The green bar marks the region of protofilamental interface. Residues forming the steric zipper in the interface are highlighted in blue. Residues forming π-stacking and hydrophobic interactions flanking the steric zipper are highlighted in orange. **b**, Top view of the hnRNPA1 LC fibril structure. Side chains that are involved in the inter-protofilamental interactions are shown in spheres. Among them, the polar side chains in the steric zipper are colored in blue; the Phe and Pro side chains are colored in orange. The two protofilaments are colored in magenta and gray, respectively. The framed regions are zoomed in and shown in top and side views in (**c-e**).

### PY-NLS is the main component of the fibril core

PY-NLS is structurally disordered in free hnRNPA1. In the complex with Kapβ2, residues 263–289 of PY-NLS forms an extended conformation to line in the C-terminal arch of Kapβ2^8^ (**Fig. 4a and 4b**). Strikingly, upon amyloid fibril formation, this same region forms the main part of FC (**Fig. 4a and 4c**). In contrast to the overall extended conformation in the complex with Kapβ2, PY-NLS forms a more compact structure in the fibril **(Fig. 4d)**. F263-F273 further extends to a β strand, referring to which the rest of PY-NLS curls into a serpentine fold **(Fig. 4d)**. The distance between the backbones of the terminal residues reduces from 42.0 Å to 22.7 Å.

**Figure 4.**
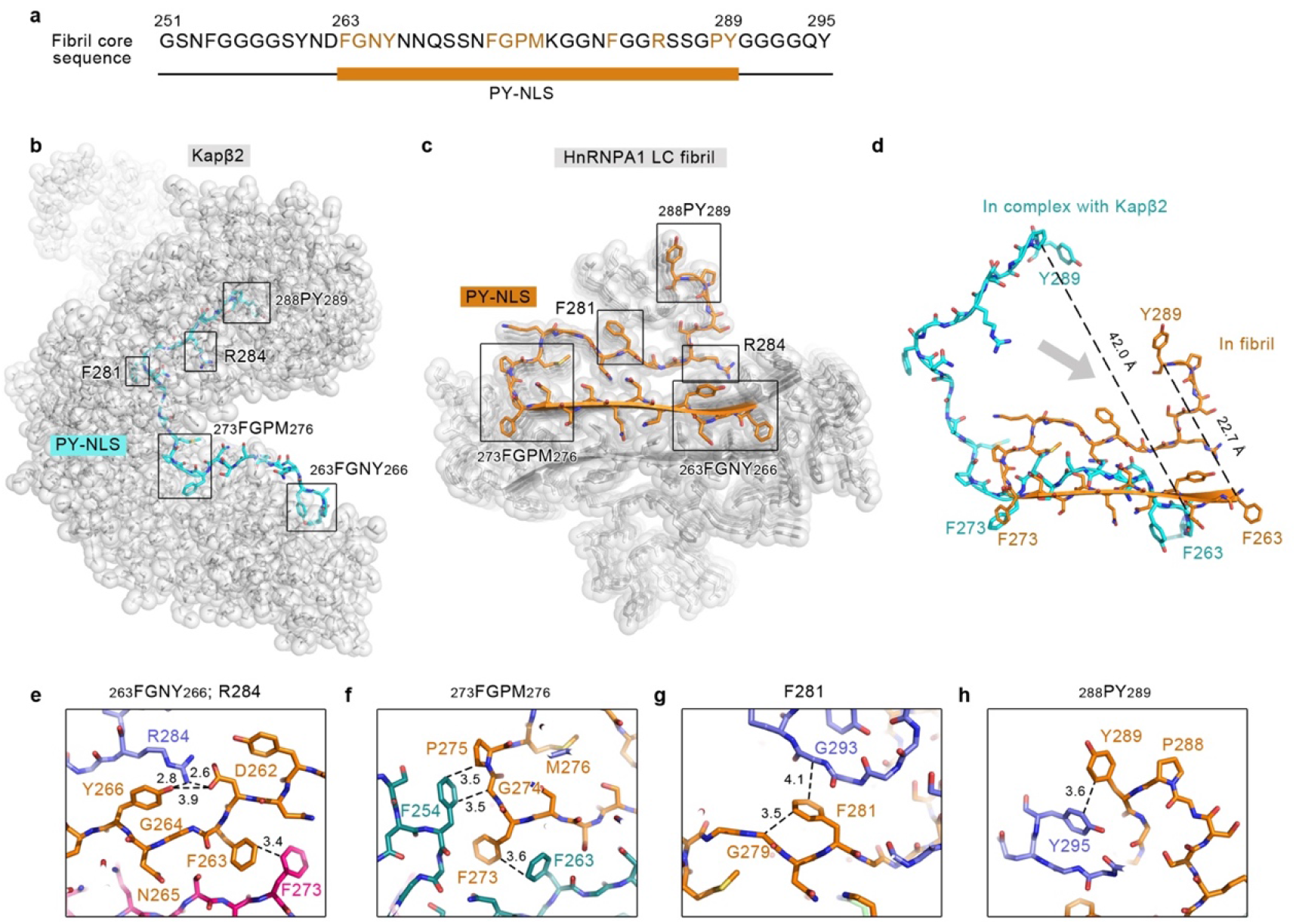
Structures of PY-NLS in complex with Kapβ2 and in the hnRNPA1 LC fibril. **a**, The primary sequence of the hnRNPA1 FC. The orange bar marks the region of PY-NLS. Residues interacting with Kapβ2 are highlighted in orange. **b**, Complex structure of PY-NLS and Kapβ2 (PDB ID: 2H4M). PY-NLS is shown in sticks and colored in cyan. Kapβ2 is shown in gray spheres. Regions of PY-NLS interacting with Kapβ2 are framed and labeled. **c**, Top view of hnRNPA1 LC fibril structure. 6 layers are shown. PY-NLS in one subunit is highlighted in orange. The rest of the structure is shown in gray spheres. The same framed regions of PY-NLS in (**b**) are framed and labeled. **d**, Overlay of PY-NLS structures in complex with Kapβ2 and in fibril. **e-h**, Zoom-in views of the framed regions in (**c**). HnRNPA1 LC subunits are colored differently. Distances between side chains are indicated in Å.

Moreover, PY-NLS residues that form extensive interactions with Kapβ2 also form important intra- and inter-molecular interactions to stabilize the amyloid fibril structure. In detail, segments _263_FGNY_266_ and _273_FGPM_276_ that contact hydrophobic patches on Kapβ2 are essential for the formation of the protofilemental interface (**Fig. 4b, 4c, 4e and 4f**). In addition to the inter-protofilamental π-stacking between F263 and F273 (**Fig. 3d and 4e**), and hydrophobic interaction between F254 and P275 (**Fig. 3e and 4f**) mentioned above, Y266 is involved in the inter-molecular electrostatic interaction between D262 and R284, and forms π-cation interaction with R284 **(Fig. 4e)**. This triad of D262, Y266 and R284 provides major forces to lock coil 2 with the β strand (**Supplementary Fig. 4**). R284 also contributes to the binding of PY-NLS with Kapβ2 by forming salt links with the acidic interface of Kapβ2^8^(**Fig. 4b**). Upon Kapβ2 binding, F281 and _288_PY_289_ also bind hydrophobic surfaces of Kapβ2. PY has been suggested as a consensus motif in various NLSs including hnRNP D, F, M, FUS, HuR, TAP, and PQBP-1, mutation of which validates the importance of this motif in the Kapβ2 binding^8,9,27–29^. Upon fibril formation, F281 forms intra- and inter-molecular hydrophobic interactions with the main chains of G279 and G293 (**Fig. 4g**); F289 of the PY motif forms π-stacking with Y295 (**Fig. 4h**). These interactions provide important forces for the folding of coil 2 (**Supplementary Fig. 4**).

Therefore, these structures demonstrate that hnRNPA1 uses the same sequence of PY-NLS for Kapβ2 binding and amyloid aggregation. Thus, the binding of PY-NLS with Kapβ2 can directly prevent it from driving the amyloid fibril formation of hnRNPA1, which provides structural understanding for the chaperone activity of Kapβ2 in the inhibition and disaggregation of hnRNPA1 fibril formation^20^.

### Disease-causative hereditary mutations in the hnRNPA1 LC fibril structure

Several ALS- or MSP-causative familial mutations of hnRNPA1 have been identified including D262N/V, N267S, and P288S/A^15–17^. In terms of Kapβ2 binding for nucleo-cytoplasmic transport, the complex structure of PY-NLS and Kapβ2 shows the binding of residues 263-289 with Kapβ2^8^. Thus, being outside of the binding region, the D262N/V mutation is less likely to influence the cellular localization of hnRNPA1. In contrast, D262 forms salt bridge with R284 in the fibril structure (**Fig. 4e and Supplementary Fig. 4**). However, the role of this salt bridge in stabilizing the fibril structure is arguable. Previous structural study indicates that D262 is one of the key Asp residues that enable the reversibility of the hnRNAP1 fibril formed during liquid-liquid phase separation via charge repulsion^12^. D262 mutations may remove the charge repulsion and enhance the formation of pathological irreversible fibril^12,30^. Without D262 mutation, wild-type hnRNPA1 is still prone to form irreversible fibrils, in which the charge repulsion of aligned D262 along the fibril is neutralized by the aligned R284 to stabilize the fibril (**Fig. 4e and Supplementary Fig. 5a**). In the context of D262N/V mutation, R284 may still form π-cation interaction with Y266 to stabilize the fibril (**Fig. 4e**). Therefore, the pathology of D262N/V mutation may mainly come from the impairment of reversible fibril formation during phase separation^12^, which consequently enhances pathological irreversible fibril formation.

N267 and P288 are involved in both Kapβ2 binding and fibril formation. Upon Kapβ2 binding, N267 forms intra- and inter-molecular hydrophilic interactions with neighboring Q269 of hnRNPA1 and N803 of Kapβ2 (**Supplementary Fig. 5b**). Upon fibril formation, N267 locates at the center of the protofilamental interface, and together with N265, Q269, and S271 forms steric zipper to stabilize the fibril interface (**Fig. 3c and Supplementary Fig. 3**). Thus, N267S mutation may influence both Kapβ2 binding and fibril formation. As for the P288S/A mutation, although P288 is involved in the fibril core, its side chain does not interact with other residues except for stacking along the fibril (**Supplementary Fig. 5a**). While, P288 is essential for Kapβ2 binding^8^ (**Fig. 4b**). Thus, the P288S/A mutation may not directly influence the fibril formation of hnRNPA1. Rather, it disrupts the proper cellular localization of hnRNPA1^17^, which may lead to cytoplasmic accumulation and amyloid aggregation of hnRNPA1.

## Discussion

HnRNPA1 is one of the most abundant nuclear proteins and shuttles between nucleus and cytoplasm to regulate gene expression and RNA metabolism^2,4,5,31^. It is composed of 320 amino acids, and nearly half of the sequence at the C-terminus is of low complexity and intrinsically disordered. The LC domain underpins the nucleo-cytoplasmic transport, liquid-liquid phase separation, and amyloid aggregation of hnRNPA1^8,11,12,14,15^. Our work reveals that the PY-NLS in the LC domain of hnRNPA1 is key not only for the nucleo-cytoplasmic transport, but also for the self-association of amyloid fibril formation. Intriguingly, hnRNPA1 can form both reversible and irreversible amyloid fibrils. The former is suggested to play a functional role in the phase separation of hnRNPA1^12^, while the latter is pathological associated with neurodegeneration^11,15^. Our previous work has identified three reversible amyloid-forming core segments (so-called RACs) of hnRNPA1, which share consensus motifs of (N)D(N) and (G)F/Y(G), and form highly reversible amyloid fibrils^12^ (**Supplementary Fig. 6**). Truncation or mutation of the key residues in RACs alters the phase separation behavior and stress granule involvement of hnRNPA1^12^. Among the three RACs, RAC3 (residues 260-266) is within the the irreversible fibril core and overlaps with PY-NLS (**Fig. 5a and Supplementary Fig. 6**), which is in line with the previous result showing that truncation of RAC3, but not RAC1 and RAC2, completely eliminates the irreversible amyloid fibril formation of hnRNPA1^12^.

**Figure 5.**
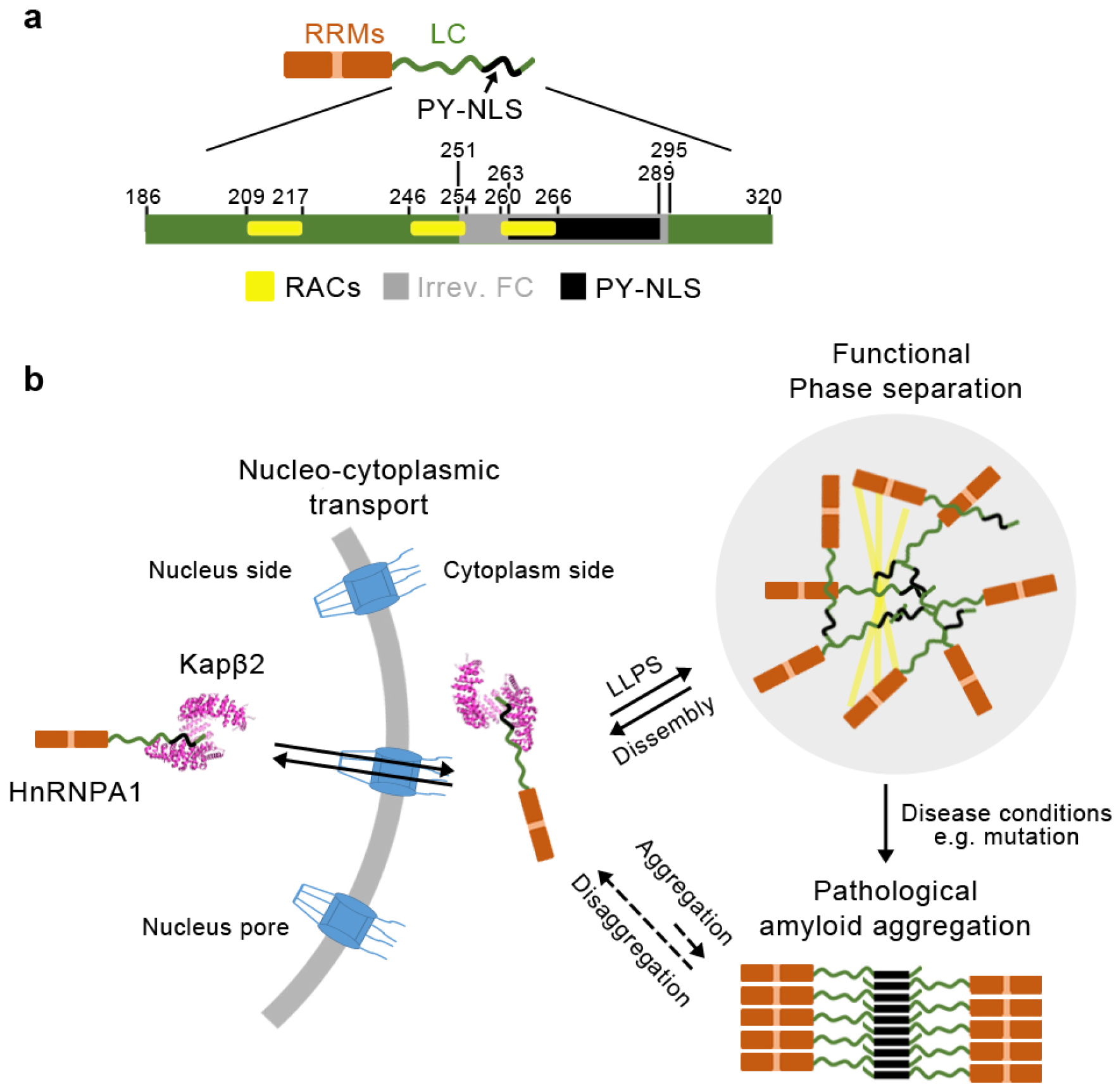
Hypothesis of hnRNPA1 forming different structures to adapt to different function. **a**, Domain organization of hnRNPA1. The LC domain is colored in green and zoomed in. Segment 251-295 that is involved in the the irreversible fibril core is colored in gray. The PY-NLS (segment 263-289) is highlighted in black. The reversible amyloid-forming core segments (RACs) are highlighted in yellow. **b**, Schematic illustration of the different structural states of hnRNPA1. HnRNPA1 monomer forms complex with Kapβ2 via PY-NLS. The LC domain mediates self-association of hnRNPA1 to undergo liquid-liquid phase separation (LLPS) via multivalent and reversible amyloid-like interactions. Under disease conditions, hnRNPA1 further assembles into highly-ordered amyloid fibrils via PY-NLS.

Combining previous and the current work, we hypothesize that hnRNPA1 forms different structures via its LC domain to adapt to different functions. Under physiological conditions, hnRNPA1 shuttles between nucleus and cytoplasm via its PY-NLS forming an extended structure lining in Kapβ2 (**Fig. 5b**). In response to cellular stimuli, the LC domain self-associates via multivalent interactions^11,14^ and reversible amyloid-like interactions^12,14,30^ to drive the liquid-liquid phase separation and incorporation of hnRNPA1 into stress granules (**Fig. 5b**). Note that the intrinsically disordered sequences of other RNA-binding proteins, e.g. TDP43, may also associate with hnRNPA1 for phase separation^13^. The involvement of the LC domain in either the Kapβ2 binding or self-association in the dynamic liquid-liquid phase separation may protect hnRNPA1 from irreversible aggregation. However, under pathological conditions such as hereditary mutations, decreased binding affinity with Kapβ2 or disruption of phase separation dynamics may expose hnRNPA1 LC, particularly the PY-NLS, and thus lead to the pathological amyloid fibril formation and neurodegeneration (**Fig. 5b**).

## Methods

### Expression and purification of hnRNPA1 LC

The gene encoding hnRNPA1 LC (residues 186-320) was inserted into pET32M vector with a Trx1 tag. HnRNPA1 LC was overexpressed in *E. coli* BL21 (DE3) at 25 ℃ overnight after induction by 0.4 mM isopropyl-1-thio-D-galactopyranoside (IPTG). Cells were harvested by centrifugation and lysed in the lysis buffer (50 mM Tris-HCl and 6 M guanidine hydrochloride, pH 8.0) at room temperature. Cell lysate was sonicated for 30 min, followed by centrifugation at 30,966 g for 45 min at 4 °C. The supernatant was loaded into Ni column (HisTrap FF, GE Healthcare), and HnRNPA1 LC was eluted with the elution buffer containing 50 mM Tris-HCl, 6 M guanidine hydrochloride and 50 mM imidazole, pH 8.0. The protein was further purified by HPLC (Agilent 1260 Infinity, Agilent Technologies), with the elution buffer containing 20%-80% (v/v) acetonitrile. After freeze-dried, the purified hnRNPA1 LC was freeze-dried and resuspended with 50 mM Tris-HCl, 100 mM NaCl, pH 7.5. The Trx1 tag was cleaved by 3C enzyme with a 50:1 (protein: enzyme) molar ratio at room temperature overnight. After centrifugation at 3,724 g for 30 min at room temperature, the precipitate was dissolved by the lysis buffer. The hnRNPA1 LC protein without tag was obtained by additional purification by HPLC, and freeze-dried.

### Fibril formation

25 μM hnRNPA1 LC in the buffer containing 50 mM Tris-HCl, 100 mM NaCl, pH 7.5 was incubated at 16 °C for 2 days without agitation. The fibrils were harvested and dialyzed into 10 mM acetate buffer (pH 5.0) overnight for cryoEM sample preparation.

### CryoEM data collection

HnRNPA1 LC fibrils were loaded onto glow-discharged copper Quantifoil grids (R1.2/1.3, 300 mesh) and then plunge-frozen in liquid ethane precooled by liquid nitrogen using Vitrobot Mark IV(FEI). Micrographs were captured using a Gatan K2 Summit camera on a Titan Krios transmission electron microscope (FEI) operated at 300 kV with a GIF Quantum energy filter (slit width 20 eV), and collected in dose fractionation mode (32 frames per micrograph) by Serial EM software. A magnification of 130,000× with physical pixel size 0.52 Å in super-resolution mode was applied. More details are shown in **Table1**.

### Imaging processing

Motion correction implement of MotionCorr2 was used to correct beam induced motion of movie frames with dose weighting implement^32^. All 32 frames per micrograph were aligned with 1.356 Å physical pixel size. CTFFIND4.1.8 was then used to estimate the contrast transfer function (CTF)^33^. All filaments were picked manually using manual picking method of RELION3.0^21^.

### Helical reconstruction

Helical reconstruction was performed by RELION3.0^21^. A box size of 288 pixels with an inter-box distance of 30 Å was used in extraction method and 801,707 segments were obtained for subsequent reference-free 2D classification. A staggered arrangement was observed in 2D classification results suggesting a pseudo-2_1_ symmetry. An initial helical rise of 2.4 Å and helical twist of 179.01° calculated by the apparent crossover distance of filaments were set in 3D classification. 259,615 segments selected after 2D classification were used to construct an initial 3D model de novo. The initial 3D model low-pass filtered to 60 Å was then applied as reference map in 3D classification. Local search of symmetry to optimize of helical twist and rise was not carried out until separation of β-strands along the helical axis was shown in 3D classification. Several rounds of 3D classification with K=3 was performed to remove heterogeneous segments. Optimized parameters and selected segments were applied for 3D refinement and final reconstruction with appropriate reference. Post-processing with a soft-edge solvent mask in 20 % central Z length was performed to sharpen final reconstruction. Fourier shell correlations at FSC = 0.143 criteria were used to estimate resolution of map. A soft spherical mask without masking a 3D helix was used to gain local resolution estimates.

### Atomic model building and refinement

The central region of the density map, which exhibits the highest resolution was used to build atomic model de novo in COOT^34^. The model containing 3 layers was generated and then refined by PHENIX using the real-space refinement program^35^.

### Buried surface area calculation

The buried surface area was calculated using areaimol from the CCP4 package^36^. with a probe radius of 1.4 Å. The difference between the solvent accessible surface areas of one hnRNPA1 LC chain alone and within the 5-layer fibril structure represents half of the reported buried area.

## Data availability

Density maps of the hnRNPA1 LC fibril are available through EMDB with entry code: EMD-30235. The structural model was deposited in the Protein Data Bank with entry code: 7BX7. Structural model of Kapβ2 used in this study is available in the Protein Data Bank with entry codes: 2H4M.

## Acknowledgements

This work was supported by the Major State Basic Research Development Program (2016YFA0501902 to C. Liu), the National Natural Science Foundation (NSF) of China (91853113 and 31872716 to D. Li and C. Liu), the Science and Technology Commission of Shanghai Municipality (18JC1420500 to C. Liu), Shanghai Pujiang Program (18PJ1404300 to D. Li), the “Eastern Scholar” project supported by Shanghai Municipal Education Commission (to D. Li). The Shanghai Municipal Science and Technology Major Project (2019SHZDZX02 to C. Liu). We acknowledge the support provided by the cryoEM facility of ShanghaiTech University.

## Author contributions

D.L., and C.L. designed the project. K.Z., Y.S., and X.G. prepared the hnRNPA1 LC fibril samples. K.Z and X.G. performed the biochemical assays. Y.S., K.Z., and B.S. performed the cryo-EM experiments, built and refined the structure model. All the authors are involved in analyzing the data and contributed to manuscript discussion and editing. D.L. and C.L. wrote the manuscript.

